# CMOS electrochemical imaging arrays for the detection and classification of microorganisms

**DOI:** 10.1101/2021.04.21.440070

**Authors:** Christopher E. Arcadia, Kangping Hu, Slava Epstein, Meni Wanunu, Aaron Adler, Jacob K. Rosenstein

## Abstract

Microorganisms account for most of the biodiversity on earth. Yet while there are increasingly powerful tools for studying microbial genetic diversity, there are fewer tools for studying microorganisms in their natural environments. In this paper, we present recent advances in CMOS electrochemical imaging arrays for detecting and classifying microorganisms. These microscale sensing platforms can provide non-optical measurements of cell geometries, behaviors, and metabolic markers. We review integrated electronic sensors appropriate for monitoring microbial growth, and present measurements of single-celled algae using a CMOS sensor array with thousands of active pixels. Integrated electrochemical imaging can contribute to improved medical diagnostics and environmental monitoring, as well as discoveries of new microbial populations.

## I. Introduction

Decades of exponential improvements in semiconductor technology scaling have brought us to a point where modern transistors approach the dimensions of large biomolecules, and whole circuit blocks may occupy footprints smaller than the area of a single cell [1]. These small feature sizes offer interesting possibilities for developing sensors with spatial resolution at the scale of single cells. Indeed, there are many examples of using semiconductor sensor arrays to record transient biopotential spikes from single excitable cells [2], and capacitance sensing arrays have been used to monitor cultured cell proliferation [3]. However, relatively fewer examples exist which interface CMOS electronics with microorganisms rather than mammalian cells and tissues.

Microorganisms include all living things with microscopic dimensions. A majority of microorganisms are unicellular, and this broad category includes species from all branches of life, including archaea, protists, bacteria, and fungi. Single cells may range in diameter from 0.1 *µ*m to 100 *µ*m, although unicellular organisms are often found in communities and their morphologies are highly dependent on their environment. In terms of sheer numbers of unique species, microbes represent the majority of the diversity of life on earth, and billions of species of microorganisms may remain undiscovered [4].

Developing integrated sensors for tracking diverse microorganisms could be especially useful for distributed environmental monitoring [5], where sensing living organisms can provide important information which is complementary to metagenomic studies. The diversity and abundance of microorganisms in field collected samples can serve as useful metrics for assessing the ecological health of soils, rivers, and oceans [6]. These types of studies are increasingly important in monitoring the effects of global warming on ecosystems.

In this paper we will summarize recent advances in CMOS electrochemical imaging for detecting and classifying microorganisms, and present new experimental recordings of single algae cells using an electrochemical impedance spectroscopy (EIS) sensor array [7]. Integrated electrochemical microbial sensing has the potential to lead to improved medical diagnostics, high throughput phenotyping, and *in situ* monitoring of microorganisms in natural environments.

## II. From Standard Microscopy to Contact Imaging

Some relevant examples of microscale imaging arrays have been adapted from standard optical microscopy. It is possible to image samples without lenses [8], especially when the distance between the object and the photodiode array becomes very small. Many modern image sensors have pixel sizes of only 1-2 *µ*m. In addition to resolving objects larger than single pixels, lens-free imaging can support computational imaging modes when combined with structured illumination [9], [10]. However, there are limits to what can be measured optically, and other sensing modalities may offer complementary forms of information.

## III. Electrochemical imaging

When a sample can be placed in direct contact with a sensor, it becomes possible to consider interrogating it electrochemically instead of optically. Electrochemical sensor arrays can be miniaturized and integrated, offering both challenges and opportunities [11]. By their nature, microscale electrochemical measurements are often noisy and prone to drift and fouling, but they can also be quite fast and low cost. In some applications, non-optical imaging could avoid complications with dyes and photobleaching, and monitor organisms independent of lighting conditions.

### A. pH Imaging

One type of chemical sensing which is readily achieved with semiconductor technology is pH sensing using ion sensitive field effect transistors (ISFETs). An ISFET is a transistor whose floating gate is sensitive to the surface potential of an exposed electrode [12]. By selecting an electrode material with pH-sensitive surface charge groups, the transistor’s inversion charge can be made a function of the pH. ISFETs can be constructed using many common oxides and nitrides, including SiO_2_, SiN, Al_2_O_3_, HfO_2_, and Ta_2_O_5_.

Integrated ISFET arrays have been designed at very large scales, especially for highly parallel DNA sequencing [13], [14]. ISFET arrays could also be interesting for monitoring microorganisms, as pH can serve as a proxy for pCO_2_ [15] and other measures of metabolism [16]. When combined with an ion-selective membrane, an ISFET may also be designed to detect other ions, such as potassium [17] or calcium [18].

### B. Reduction–Oxidation Imaging

In addition to sensing bulk chemical properties such as pH, other electrochemically active species can also provide information about a microorganism. Some researchers have developed systems which can produce electrochemical images of electroactive metabolites as they are released by tissue samples [19] or bacterial biofilms [20], [21]. Cells on microelectrode arrays can also measurably affect the transport of bulk redox species to electrodes [22], [23].

### C. Impedance and Capacitive Imaging

Electrochemical impedance spectroscopy (EIS) is traditionally used to characterize electrode surfaces, for applications including corrosion monitoring or characterizing the fouling of implanted electrodes [24], [25].

It is also possible to use EIS to monitor the growth of adherent cells on surfaces [3], [26]–[29], or to estimate the surface coverage of bacterial biofilms [30]. At larger dimensions, this approach has been commercialized in impedance-based cell culture monitoring systems, which use millimeter-scale interdigitated electrodes [26].

These examples use EIS for monitoring cells in extremely close contact with the electrode because mobile dissolved ions in the buffer form an electrostatic double layer at the surface with very high charge density [31]. This Debye layer screens the electric field from penetrating more than a few nanometers into a sample. However, the relaxation time of the screening layer is on the order of 1 *µ*s, and at MHz frequencies the electric field can extend farther into a sample [32]–[34]. Radiofrequency EIS offers opportunities for imaging thicker samples or non-adherent cells. Operating at higher frequencies also makes an impedance measurement less dependent on the exact surface charge, which is prone to drift.

Similar to optical phase contrast imaging [35], EIS imaging can observe changes in local dielectric properties, such as water displaced by a cell crowded with lipids, proteins, and other molecules. Under some conditions, impedance imaging can detect and localize conductivity changes more than 100 *µ*m from the electrode surface [36].

## IV. Impedance Imaging of Single-Celled Algae

To illustrate some of the types of microbial imaging that can be done using electrochemical sensors, we measured a variety of unicellular algae using a high-frequency EIS CMOS sensor array. The design details of the integrated circuit are described in [7] and its operating principle is similar to those described elsewhere [33], [34], [37]. Briefly, the sensor contains a grid of electrodes with a pitch of approximately 10 *µ*m. Each electrode can be rapidly charged and discharged between two bias voltages, while the net current is measured (Fig. 2c). The switching frequency can operate as fast as 100 MHz, and the charge transferred per cycle is a function of the interfacial capacitance (alternatively expressed in terms of impedance) between the metal electrode and the wet sample on its surface. A silver/silver-chloride reference electrode maintains the solution at a constant potential, although the measurement is typically not sensitive to the exact DC solution bias.

**Fig. 1.**
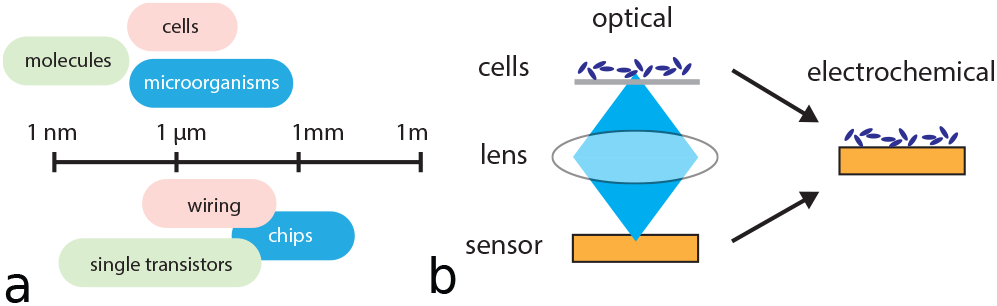
Overview. (a) The dimensions of modern integrated circuits overlap with the scales of single cells. (b) Bringing samples into contact with semi-conductor chips can enable spatially-resolved electrochemical interrogation of microorganisms.

**Fig. 2.**
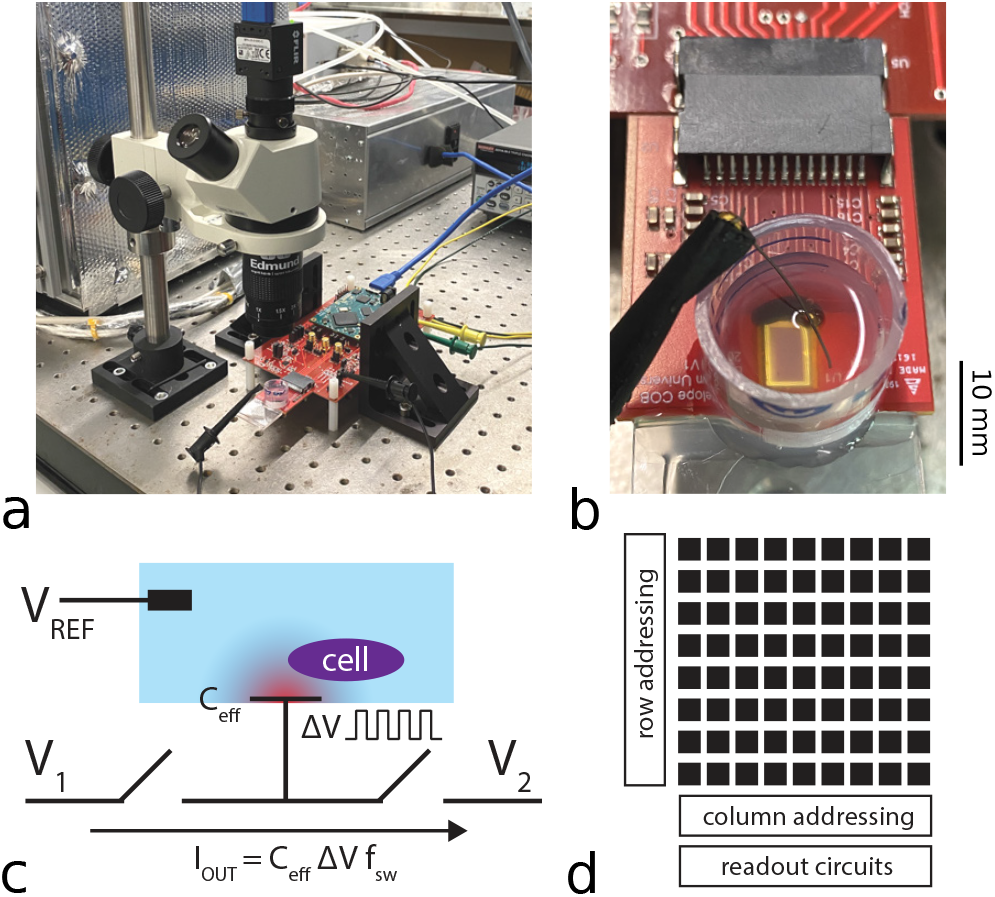
Electrochemical imaging biosensor array. (a) An integrated CMOS impedance sensor array [7] is attached to a small module which plugs into a custom data acquisition circuit board. The sensor is positioned under an inspection microscope during experiments. (b) The sensor is assembled with a simple fluid chamber and a silver/silver-chloride counter-electrode. (c) A simplified circuit diagram of the impedance sensing pixel. Non-overlapping clocks rapidly charge and discharge the electrode’s interfacial capacitance, while the total switched pixel current is measured. (d) Arrays of these sensor pixels can support spatially-resolved impedance imaging at frequencies up to 100 MHz.

After assembling a small open fluid chamber around the sensor, we arranged the system under an inspection microscope (Fig. 2), for simultaneous optical and electrochemical visualization.

Figure 3 shows measurements of filamentous freshwater green algae dispersed onto the surface of the sensor. There is strong correlation between the optical and impedance images, and the quality of the impedance image improves significantly at higher frequencies. Despite the improved sensing depth at radio frequencies, not all of the algae cells are detected in the impedance image. The rectangular cells have widths on the order of 20 *µ*m, and they do not all lie flat on the sensor surface.

**Fig. 3.**
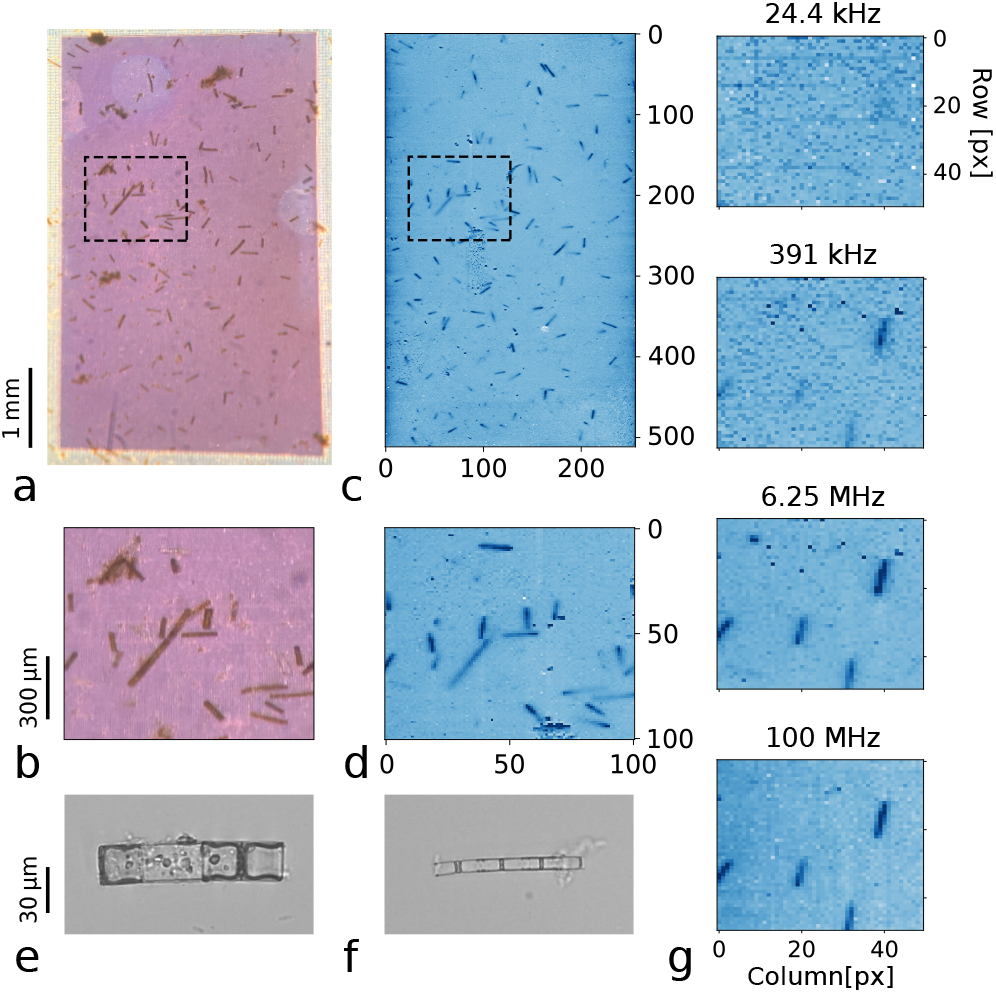
EIS imaging of freshwater green algae. (a) & (b) Images of green algae dispersed onto the surface of a CMOS sensor array. (c) & (d) EIS measurements of the algae cells. (e) & (f) Optical images of the algae cells taken with a bright-field transmitted light inverted microscope. (g) The EIS image contrast is a strong function of the switching frequency, illustrating the benefit of operating at high frequency to overcome Debye screening.

In Figure 4 we see measurements of *Cosmarium turpinii*, a freshwater algae with a cell diameter of approximately 50 *µ*m and a notable constriction in the middle. Individual cells are clearly detected by CMOS impedance imaging, although the subcellular structure is only sometimes observable. Cells with consistent and distinct shapes offer interesting test cases for evaluating the spatial resolution of CMOS biosensors.

**Fig. 4.**
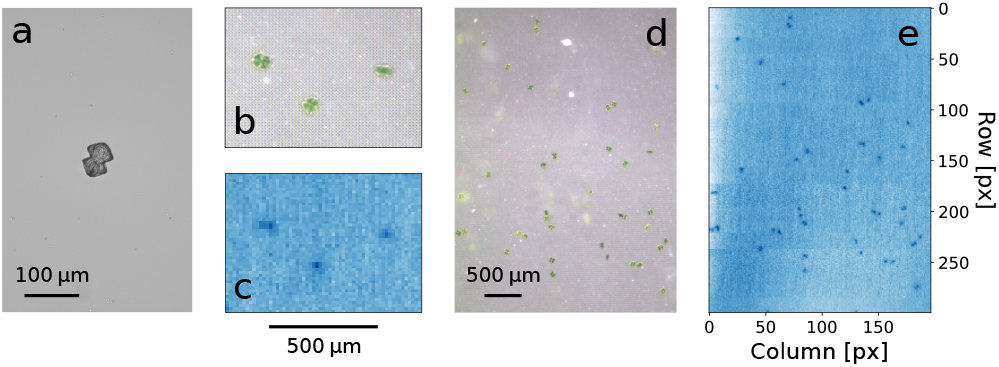
*Cosmarium turpinii* algae. (a) Microscope image of a single cosmarium turpinii cell. (b) Optical image of three cells on the array. (c) Corresponding EIS measurement. (d) A collection of cosmarium cells on the CMOS sensor. (e) Corresponding EIS image captured by the sensor.

Figure 5 shows images of *Closterium acerosum*, another type of freshwater green algae which has crescent-shaped cells with tapered ends. These larger cells are very clearly resolved, though on occasion we can see some loss of contrast as part of the cell appears to extend farther from the sensor surface.

Smaller microalgae are more challenging to detect. Figure 6 shows measurements of *Cyclotella* sp., which are round and flat marine diatoms with diameters close to the size of the 10 *µ*m sensor pixels. Some of these cells are clearly detected, but they appear primarily as single points in the EIS image.

**Fig. 5.**
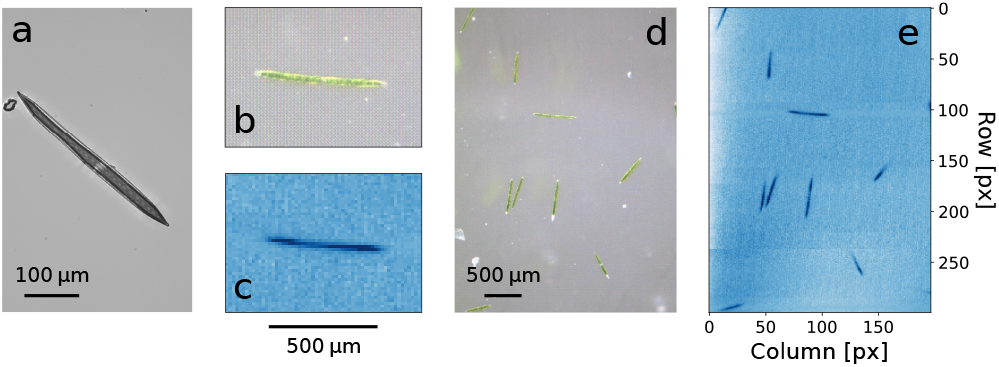
*Closterium acerosum* algae. (a) Microscope image of a single closterium cell. (b) Optical image of a closterium cell on the array. (c) Corresponding EIS measurement. (d) A collection of closterium cells on the CMOS sensor. (e) Corresponding EIS image captured by the sensor.

**Fig. 6.**
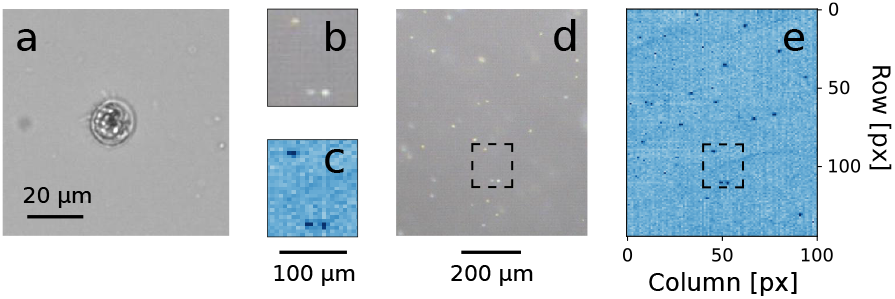
*Cyclotella* sp. microalgae. (a) Optical image of a cyclotella cell. (b) A magnified area of the optical image. (c) Corresponding EIS measurement. (d) Cyclotella dispersed on a CMOS sensor. (e) An EIS image of cyclotella cells. (In this example, the solution was mixed between optical and EIS imaging, displacing some of the cells. The dashed boxes highlight a group of cyclotella that remained in position.)

Impedance images can be resolved temporally as well as spatially [30]. Closterium (Fig. 5) can move by secreting mucilage to push away from objects in their environment. Figure 7 shows snapshots from a 1.5 hour impedance time-lapse movie, in which single closterium cells can be seen moving laterally across the sensor and vertically in and out of the sensing volume. Visualizing the kinetics of cellular growth and motion offer another opportunity for characterizing and identifying microorganisms.

**Fig. 7.**
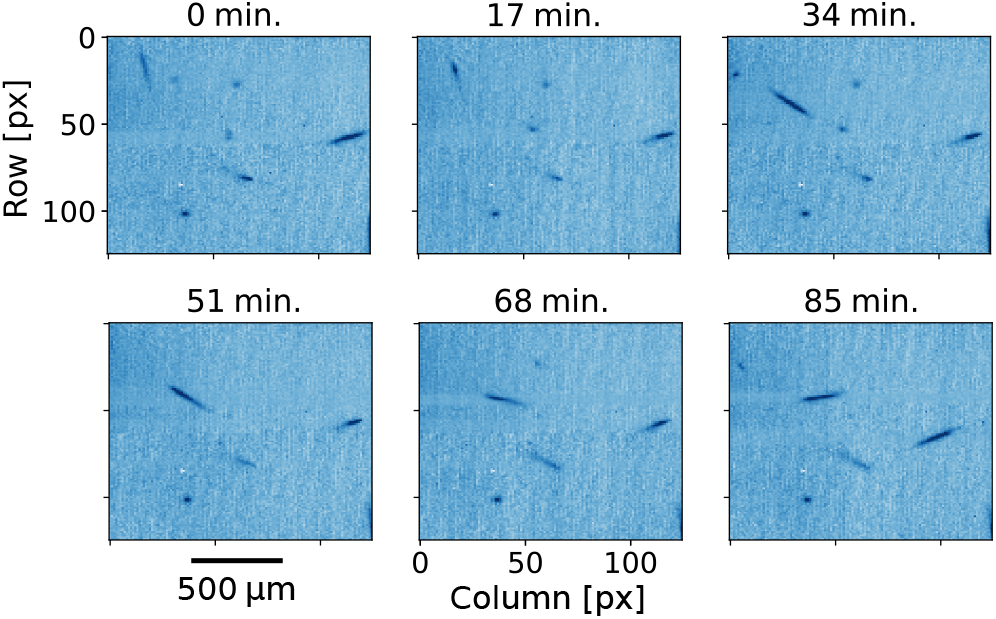
Single cells in motion. Snapshots from a time lapse recording of EIS images showing several closterium cells moving around on the surface, over the course of 85 minutes.

Clearly any natural environment will present mixtures of species, rather than pure cultures. Figure 8 shows a mixture of cosmarium and closterium algae cells dispersed onto an EIS array. Even with this simple mixture, we can begin to appreciate the challenge of not merely detecting cells, but also classifying them.

**Fig. 8.**
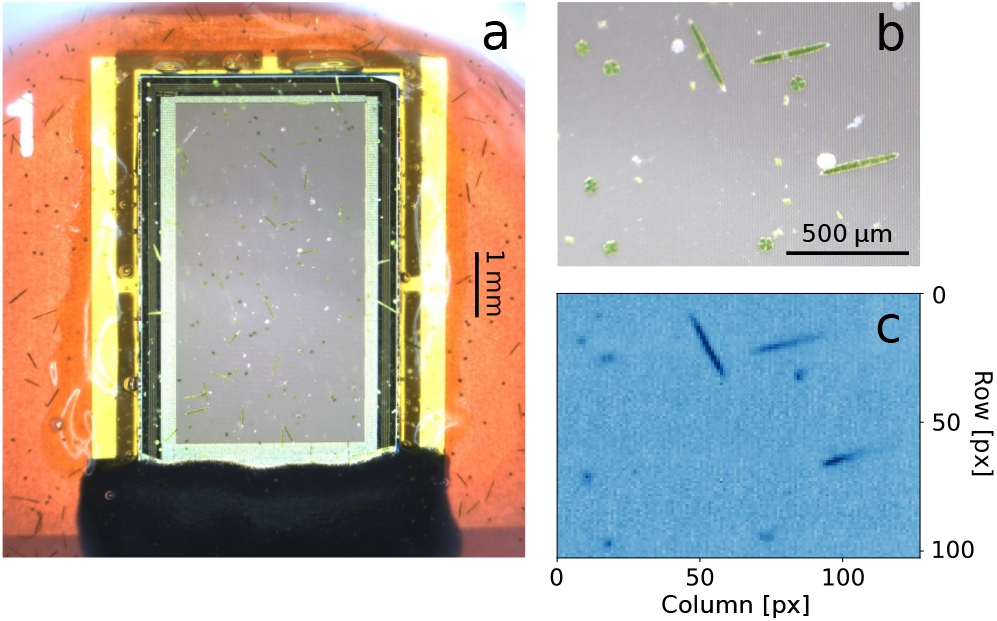
Mixed algae sample. (a) Image of a mixture of *Cosmarium turpinii* and *Closterium acerosum* cells dispersed on the CMOS sensor. (b) A magnified microscope image of the mixed algae sample on the sensor. (c) Measured EIS image of the mixed sample.

In Figure 9, we show an example from a dataset in which cells were segmented from the background, and a classifier was trained on images of pure cosmarium and closterium cells. The classification was based on two simple shape metrics, cell area and aspect ratio. This classifier was then applied to a mixed sample, where it was able to successfully label a large majority of the cells. Most of the errors occurred when closterium cells were tilted out of the sensing plane and mistaken for smaller cosmarium cells. Automated phenotype classification from microscopy data is an exciting and active area of research [38], [39], and as the spatial resolution of electrochemical images improves, there will be more opportunities for applying machine learning to these rich datasets.

**Fig. 9.**
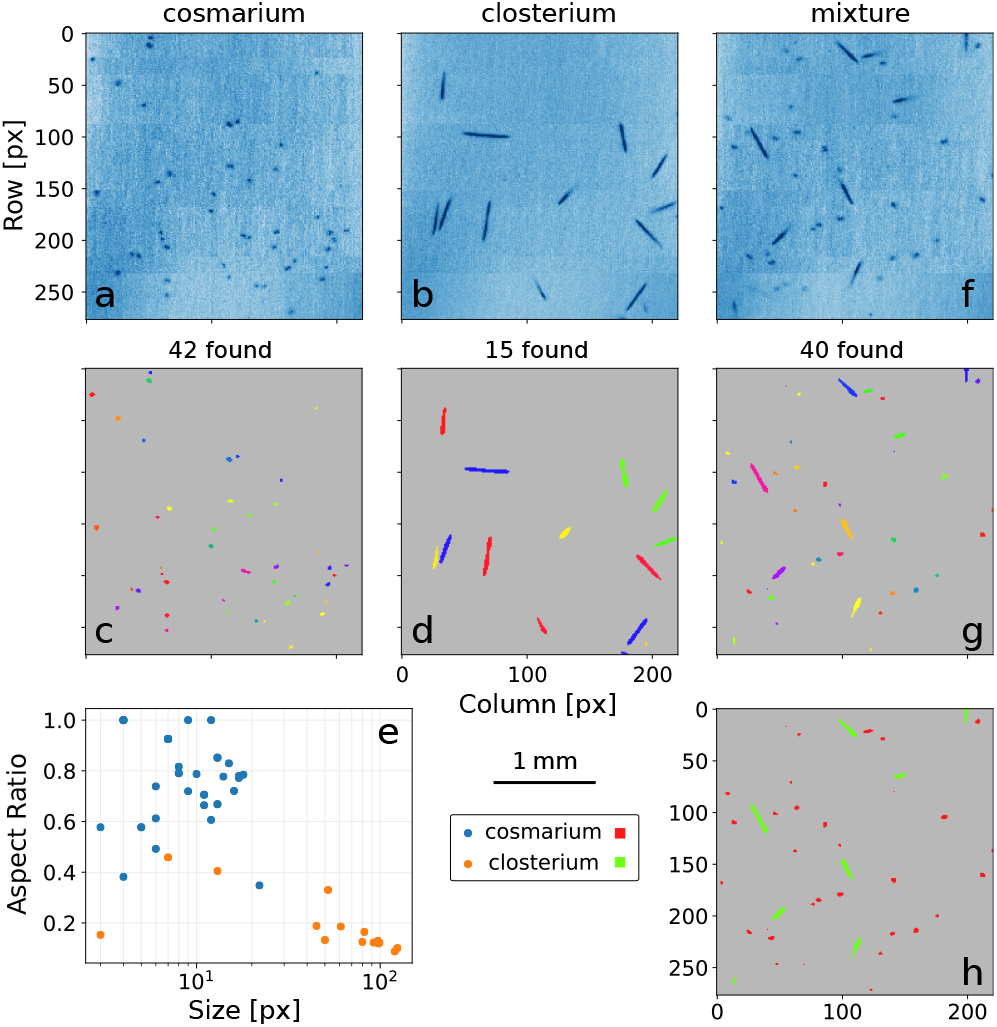
Classifying a mixed algae population. (a) & (b) Monocultured algae samples were measured independently. (c) & (d) Thresholded and segmented images of each type of algae. (e) The cells were characterized by size and aspect ratio using a multi-class Gaussian process model. (f) A measured sample containing a mixture of algae species. (g) Segmented image of the mixed sample. (h) Cells from the mixed sample were classified and labeled using the trained model.

## V. Conclusion

In this paper we have presented high-frequency impedance images of single algae cells, recorded on an integrated CMOS electrochemical sensor array. These results highlight the continued opportunity to use modern semiconductors for miniaturized biosensors and bioimaging platforms. CMOS electrochemical sensor arrays offer not only low cost and massive parallelism, but also opportunities for spatially resolved measurements of biological samples at the scale of single cells.

The compact profile of lens-free imaging arrays may open up applications for monitoring microbes in environments where conventional microscopes would be cumbersome or infeasible. The absence of optical illumination makes electrochemical sensors especially well suited for monitoring photosensitive organisms.

Limitations of CMOS electrochemical sensor arrays include their finite out-of-plane sensing distance, which generally limits their application to relatively thin samples. Data acquisition can also be fairly slow, with familiar tradeoffs between resolution, pixel count, and frame rate. Fortunately, imaging cell growth generally does not require high frame rates.

A tremendous amount of information is contained in the geometry and spatial structure of cells. If we can miniaturize sensor arrays to achieve spatial resolution comparable to optical microscopy, then we can detect and classify some organisms by label-free general-purpose electrochemical imaging.

## Supplementary Methods

### 1. Instrumentation

Electrochemical images were acquired using a custom CMOS sensor, which is described in Hu et al. 2021 [1]. The sensor supports electrochemical impedance spectroscopy (EIS), pH imaging, and optical imaging, but only the EIS mode was used here. Impedance images were taken using a switching frequency of 100 MHz, unless otherwise stated. The resulting images were processed and analyzed with Python within Anaconda3, using scikit-image [2] and scikit-learn [3]. Reference optical images of algal cells (Fig. 3e-f, Fig. 4a, Fig. 5a, Fig. 6a) were taken using a Nikon TI-U inverted microscope (bright-field, 20 *×* magnification). Optical images of the cells on the CMOS sensor (Fig. 4b & d, Fig. 5b & d, Fig. 6b & d, Fig. 8a & b) were taken using the inspection microscope shown in Fig. 2a (Edmund Optics #55-150 dual tube body, 0.75 - 3 *×* magnification, FLIR BFS-U3- 51S5C-C camera).

### 2. Living Specimen

Live algae were purchased from Carolina Biological Supply Company (Burlington, NC, USA). The unidentified filamentous freshwater green algae were taken from a field-collected mixed culture (Item #151287). The *Cosmarium* (Item #152140), *Closterium* (Item #152115), and *Cyclotella* (Item #153020) cells were obtained as unialgal cultures. For classification tests, we used these samples to prepare a mixture of *Cosmarium* and *Closterium* cells.

### 3. Experimental Setup

Before each experiment, a small fluid chamber (Fig. 2b) is fashioned out of a centrifuge tube section and bonded to the surface of the sensor using silicone elastomer. The chamber is then filled with phosphate-buffered saline (0.1 M potassium phosphate, 1 M potassium chloride). An Ag/AgCl wire, serving as a reference electrode, is placed into the solution. The entire setup is placed under an inspection microscope and algae samples are pipetted into the chamber. After the cells settle onto the sensor surface, optical images are taken for reference and electrochemical imaging is performed. Each 512 *×* 256 impedance image takes approximately one minute to acquire.

## Notes

This work was supported in part by a grant from the Defense Advanced Research Projects Agency (DARPA). The views, opinions and/or findings expressed are those of the authors and should not be interpreted as representing the official views or policies of the Department of Defense or the U.S. Government. This work was also supported in part by the National Science Foundation under Grant No. 2027108.

### Competing Interest Statement

The authors have declared no competing interest.

